# Human TLR4 and noncanonical inflammasome differ in their ability to respond to distinct lipid A variants

**DOI:** 10.1101/2021.12.16.472937

**Authors:** Jasmine Alexander-Floyd, Antonia R. Bass, Erin M. Harberts, Daniel Grubaugh, Joseph D. Buxbaum, Igor E. Brodsky, Robert K. Ernst, Sunny Shin

## Abstract

Detection of Gram-negative bacterial lipid A by the extracellular sensor, MD-2/TLR4 or the intracellular inflammasome sensors, CASP4 and CASP5, induces robust inflammatory responses. The chemical structure of lipid A, specifically the phosphorylation and acylation state, varies across and within bacterial species, potentially allowing pathogens to evade or suppress host immunity. Currently, it is not clear how distinct alterations in the phosphorylation or acylation state of lipid A affect both human TLR4 and CASP4/5 activation. Using a panel of engineered lipooligosaccharides (LOS) derived from *Yersinia pestis* with defined lipid A structures that vary in their acylation or phosphorylation state, we identified that differences in phosphorylation state did not affect TLR4 or CASP4/5 activation. However, the acylation state differentially impacted TLR4 and CASP4/5 activation. Specifically, all of the examined tetra-, penta-, and hexa-acylated LOS variants activated CASP4/5-dependent responses, whereas TLR4 responded to penta- and hexa-acylated LOS but did not respond to tetra-acylated LOS or pentaacylated LOS lacking the secondary acyl chain at the 3’ position. As expected, lipid A alone was sufficient for TLR4 activation; however, human macrophages required both lipid A and the core oligosaccharide to mount a robust CASP4/5 inflammasome response. Our findings show that human TLR4 and CASP4/5 detect both shared and non-overlapping LOS/lipid A structures, which enables the innate immune system to recognize a wider range of bacterial LOS/lipid A, thereby constraining the ability of pathogens to evade innate immune detection.

## Introduction

Gram-negative bacteria are responsible for more than 30% of hospital-acquired infections in the US, making them a costly and deadly public health concern [1]. Uncontrolled Gram-negative bacterial infections can lead to detrimental outcomes including sepsis, which is an overwhelming systemic inflammatory response to an infection. If left untreated, a septic host will succumb to organ failure and ultimately death. Preclinical studies in mice successfully treated sepsis using immunomodulators that functioned by neutralizing either host inflammatory mediators or microbial products [2]. However, over a hundred clinical trials testing these immunomodulators in sepsis patients have failed. The reasons for these failures are unclear but may be due to differences between murine and human innate immune responses that play a role in responding to Gram-negative bacterial infections. Understanding further the human innate immune responses to Gram-negative bacterial pathogens will aid in identification of potential novel therapeutic targets for the treatment of Gram-negative sepsis.

Gram-negative sepsis is caused by the bacterial endotoxin, lipopolysaccharide (LPS) or lipooligosaccharide (LOS), which is the major lipid component in the outer leaflet of the outer membrane of Gram-negative bacteria. LPS is composed of a lipid A membrane anchor attached to a core oligosaccharide, which has a variable number of repeating O-antigen units attached to it. If a particular bacterial species makes lipid A with only the core attached, it is then named LOS instead of LPS. Both LOS and LPS have a lipid A moiety, which acts as a membrane anchor in the outer leaflet of the Gram-negative outer membrane. It is the lipid A subunit that activates both cell surface and cytosolic sensors, which subsequently leads to signaling events resulting in inflammatory cytokine release as well as a form of inflammatory programmed cell death termed pyroptosis. Toll-like receptor 4 (TLR4) complex is a plasma membrane-bound receptor that detects lipid A in the extracellular environment or within endosomal compartments. The lipid A/TLR4 signal transduction pathway involves binding of lipid A to the signaling cofactor myeloid differentiation 2 (MD2). Upon lipid A binding, the MD2/TLR4 complex dimerizes, leading to a conformational change in TLR4 and downstream signaling that promotes production of proinflammatory cytokines.

In addition to extracellular sensing of lipid A by TLR4, lipid A that enters the cytosol in the context of invasive bacterial pathogens or delivery by bacteria-derived outer membrane vesicles, is sensed by the cytosolic receptor caspase-11 (Casp11) in mice or the orthologs CASP4 and CASP5 in human cells [3–5]. Binding of lipid A to the caspase activation and recruitment domain (CARD) of Casp11, CASP4, or CASP5 leads to formation of a noncanonical inflammasome, resulting in autoproteolytic cleavage and caspase activation [6]. Active caspase-11, −4, and −5 subsequently cleave gasdermin-D (GSDMD), the initiator protein of pyroptosis. Importantly, both the noncanonical inflammasome and TLR4 make independent contributions to host protection in models of systemic Gram-negative infection as well as to immunopathology in models of lethal sepsis.

Lipid A is comprised of two glucosamine residues that contain hydrophobic acyl chains that vary in number, position, and length depending on the bacterial species. Also contingent on the bacterial species, lipid A possesses phosphate groups located on the 1 and/or 4’ positions of the two glucosamine residues [7]. The MD-2 coreceptor recognizes specific lipid A structures and upon binding, undergoes a conformational change which initiates the activation of TLR4, whereas intracellularly, the CARD domains of Casp11/4/5 recognize LPS to activate the inflammasome [6].

Intriguingly, pathogenic bacteria modify their acylation and phosphorylation states in response to environmental cues, suggesting that changing these vital features is important for their pathogenesis and potentially allowing these bacteria to evade immune detection or resist innate immune killing mechanisms. There are also species-specific differences in lipid A recognition as evidenced by the observations that murine and human TLR4 can differ in their responses to distinct lipid A variants [8]. For example, tetra-acylated *Yersinia pestis* lipid A evades TLR4 detection in humans while maintaining slight agonist activity for murine TLR4, whereas the hexa-acylated form robustly activates TLR4 in both mice and humans [9, 10]. Additionally, penta-acylated LPS from *Neisseria meningitidis* LpxL1 potently activates murine TLR4 but not human TLR4 [11]. Not only are there differences in how TLR4 responses to structural lipid A variants, but there are also differences in how human or murine TLR4 respond to the same lipid A.

Similar to the murine MD-2/TLR4 receptor system, the murine noncanonical inflammasome responds poorly to lipid A with a lower number of acyl chains (i.e. tetra-acylated), whereas lipid A containing higher number of acyl chains (i.e. hexa-acylated) robustly activate Casp11 for downstream pyroptosis and release of inflammatory cytokines. Interestingly, some pentaacylated lipid A variants, such as *Francisella novicida lpxF* mutant, can activate Casp11 [4], while other penta-acylated lipid A variants, like *Rhizobium galegae*, evade Casp11 detection [3]. This raises the question of whether the number of acyl chains is the sole structural feature that dictates noncanonical inflammasome activation or whether other structural features of lipid A also influence Casp11 activation. In contrast to the murine system, the human noncanonical inflammasome is activated in response to tetra-acylated lipid A, including the tetra-acylated *Francisella novicida*, as well as penta- and hexa-acylated lipid A [5]. This indicates that the human noncanonical inflammasome can be activated by lipid A with a broader range of acylation states [5]. However, whether the position of acyl chains or the number of phosphoryl groups on lipid A play a role in human noncanonical inflammasome activation has not been studied.

Here, we sought to dissect the contribution of differential acyl chain modification and glucosamine phosphorylation to the activation of the human lipid A sensing pathways. We used a comprehensive panel of lipid A structures that vary only in acylation and phosphorylation state and were isolated from acyltransferase and/or phosphatase mutants in an avirulent version of *Yersinia pestis. Y. pestis* naturally produces LOS due to a mutation that only allows for addition of a single O-antigen unit [12]. We found that differences in phosphorylation state did not affect TLR4 or CASP4/5 activation. However, the acylation state differentially impacted TLR4 and noncanonical inflammasome activation. Specifically, all examined tetra-, penta-, and hexaacylated LOS variants activated the noncanonical inflammasome, whereas TLR4 responded to penta- and hexa-acylated LOS but did not respond to tetra-acylated LOS or penta-acylated LOS lacking the secondary acyl chain at the 3’ position. Additionally, we found that while lipid A alone was sufficient for TLR4 activation, human macrophages required both lipid A and the core oligosaccharide to mount a robust noncanonical inflammasome response. In summary, our data indicate that the human noncanonical inflammasome can respond to a wider range of lipid A structures than murine Casp11, murine TLR4, and human TLR4. These findings reveal that the human cytosolic and cell surface lipid A sensing systems respond to both shared and unique structures. We expect that this broader range of lipid A structure recognition provided by human TLR4, and human CASP4/5 provides a comprehensive detection strategy that limits Gramnegative pathogen evasion of innate immune sensing.

## Results

### Structure-activity relationship of LOS with the human TLR4 signaling complex

To determine the effect of acyl-chain variation on the activation of the human TLR4 signaling complex, we treated HEK-Blue hTLR4 and THP1-Dual reporter cell lines with a series of lipid A variants that were generated in *Y. pestis (Yp)* using bacterial enzymatic combinatorial chemistry[13]. These molecules differ only in the number and position of acyl chains or phosphates in their lipid A moieties (**Supplemental Figure 1**). The HEK-Blue hTLR4 cells are co-transfected with the human TLR4, MD-2, and CD14 coreceptor genes, and both the HEK-Blue hTLR4 and THP1-Dual reporter cell lines express and secrete alkaline phosphatase downstream of NF-κB activation, allowing us to measure TLR4 activity via a colorimetric assay.

Consistent with observations made in analyses of mouse TLR4 activation in the companion paper by Harberts, Grubaugh, *et al*. [14], activation of human TLR4 in the HEK-Blue hTLR4 and THP1-Dual reporter cell lines was observed following treatment with penta-acylated *Yp*Δ*msbB* LOS lacking the secondary C12 acyl chain at the 3’ position, but not penta-acylated *Yp*Δ*lpxP* LOS lacking the secondary C16:1 acyl chain at the 2’ position (**Figure 1**). As expected, we did not observe TLR4 activation following treatment with tetra-acylated *Yp*Δ*msbB/lpxP* LOS (**Figure 1**). These data support the concept that position-dependent acyl chain additions to the base tetra-acylated bacterial lipid A structure affect human TLR4 activation. We saw similar results to LOS treatment when we instead treated our reporter cell lines with lipid A from each variant, which was derived from the same extraction lots (**Supplemental Figure 2**). These data indicate that lipid A is sufficient to stimulate TLR4 and that the core oligosaccharide moiety is not required for the TLR4-stimulating capacity of lipid A. Overall, these data indicate that the TLR4 receptor complex can be activated by lipid A and that MD-2/TLR4 is able to discriminate between penta-acylated lipid A containing a secondary acyl-chain at the 2’ position, but not the 3’ position.

**Figure 1.**
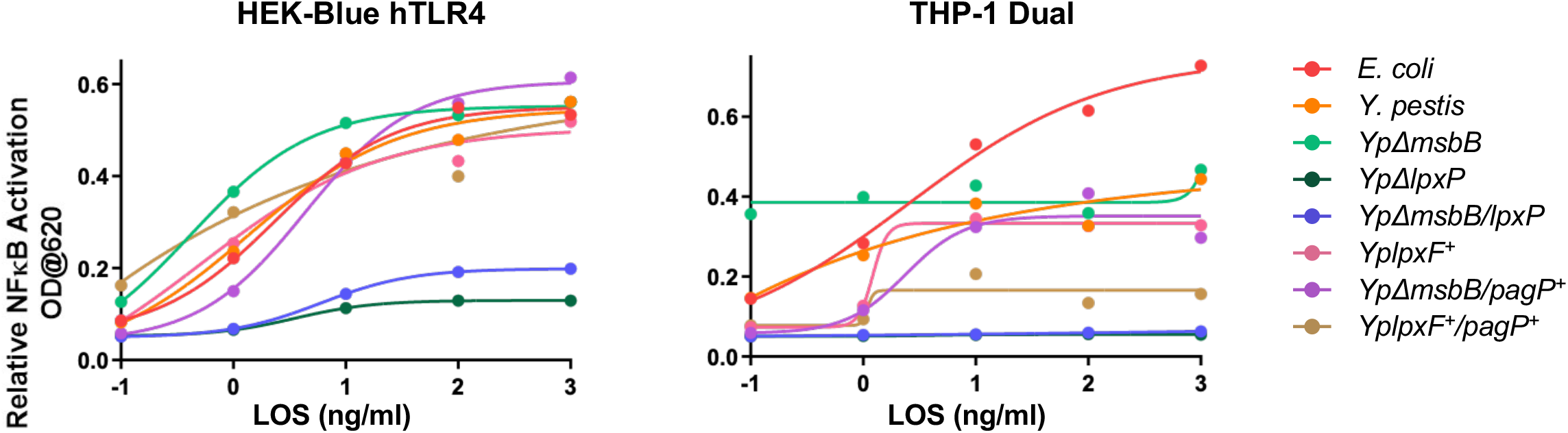
Lipid A structure determines strength of pro-inflammatory signaling downstream of TLR4. LOS structural variants were cultured with reporter cell lines that overexpress human TLR4/MD-2 (HEK-Blue hTLR4) or express endogenous levels of TLR4 (THP-1 Dual) for 18 hours. LOS variants were derived from wild type *E. coli* (red), wild type *Yp* (orange), *Yp*Δ*msbB* (light green), *Yp*Δ*lpxP* (dark green), *Yp*Δ*msbB/lpxP* (blue), *YplpxF*^+^(*pink*), *YplpxF^+^/pagP*^+^ (brown), and *Yp*Δ*msbB/pagP*^+^(*purple*). Results were graphed using GraphPad Prism v7 with a 4-parameter exponential line of best fit superimposed. Each data point is an average of biological duplicates. Representative of three experiments.

### The human noncanonical inflammasome is activated by Y. pestis LOS variants regardless of acyl chain number

Previous studies indicate that human macrophages can mount noncanonical inflammasome responses to tetra-acylated LPS [14]. However, these studies were conducted in macrophages primed with IFNγ, which may induce expression of additional host factors that can promote noncanonical inflammasome responses to LPS. IFNγ priming mimics conditions following infection when the host already has activated inflammatory pathways leading to IFNγ, not the conditions early during a primary infection. To determine how variation in acyl chain number affects activation of the human noncanonical inflammasome in the absence of IFNγ priming, we employed a well-characterized set of LOS variants derived from hexa-acylated WT *Y. pestis* (*Yp*), two penta-acylated LOS variants isolated from *Y. pestis* strains lacking the acyltransferases MsbB or LpxP that add C12 and C16:1 groups, respectively, to lipid A (*Yp*Δ*msbB and Yp*Δ*lpxP*) or the tetra-acylated LOS variant obtained from a *Y. pestis* strain lacking both MsbB and LpxP (*Yp*Δ*msbB/lpxP*)[15]. These LOS variants were transfected into primary human monocyte-derived macrophages (hMDMs) derived from 3-5 healthy human donors. We then assessed inflammasome activation by monitoring cell death via lactate dehydrogenase (LDH) release into the supernatant and IL-1β cytokine secretion by ELISA. As expected, hexa-acylated LOS derived from WT *Yp* resulted in robust cell death in a dose-dependent manner (**Figure 2A**). We also found that penta-acylated LOS from either *Yp*Δ*msbB* (**Figure 2B, 2F**) or *Yp*Δ*lpxP* (**Figure 2C, 2G**) both induced robust cell death and IL-1β secretion in hMDMs in a dose-dependent manner, at levels similar to those observed in hMDMs transfected with WT *Yp* LOS (**Figure 2A, 2E, 2I, and 2J**). Moreover, we observed that although the tetra-acylated *Yp*Δ*msbB/lpxP* LOS variant induced cell death and IL-1β secretion in a dosedependent manner, they were significantly lower compared to the levels observed in hMDMs transfected with the WT hexa-acylated variant, indicating decreased inflammasome activation (**Figure 2D, 2H, 2I, and 2J**).

**Figure 2.**
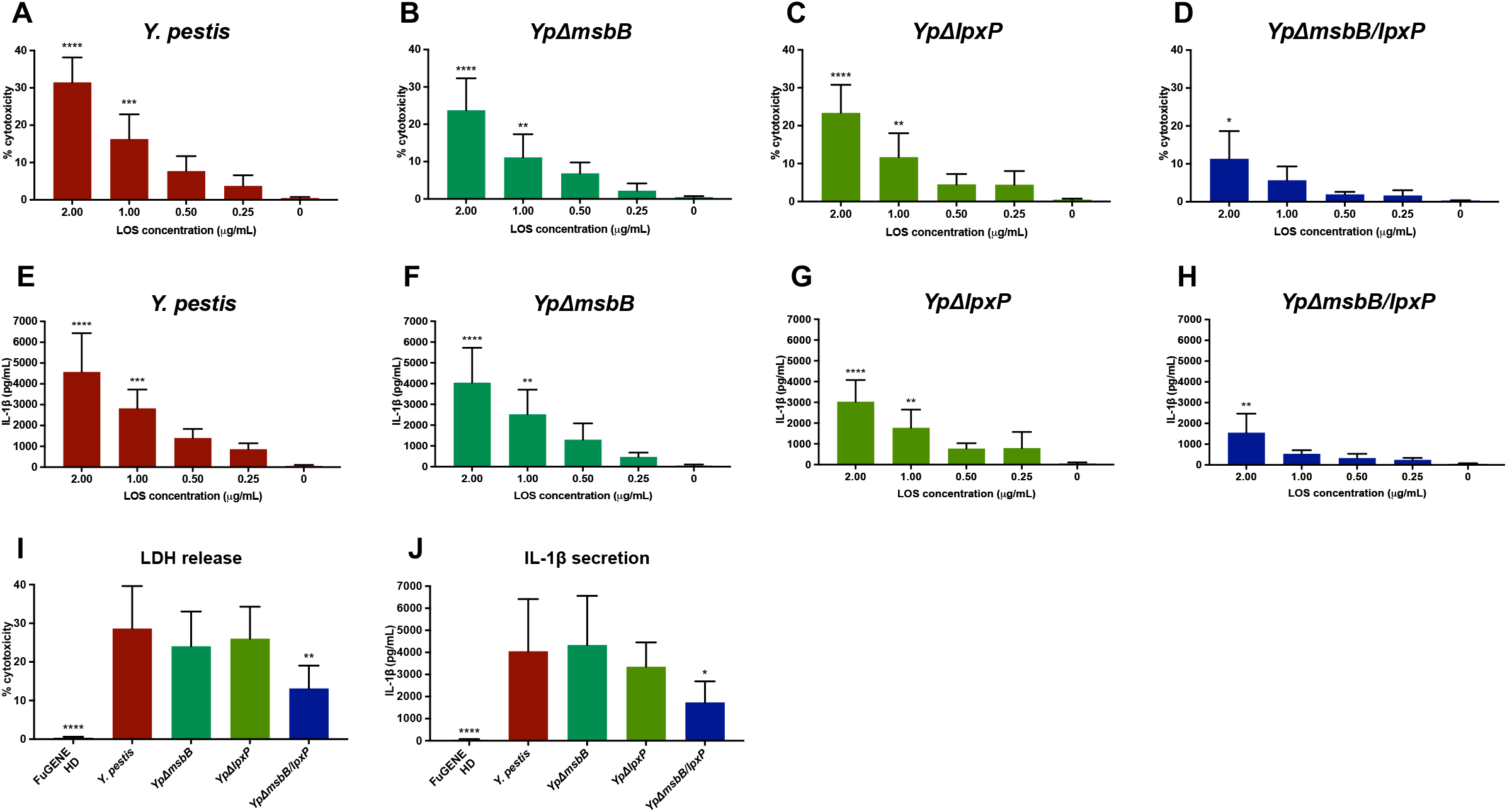
Human noncanonical inflammasome is activated by hexa-, penta-, and tetraacylated *Yersinia pestis* LOS variants. Pam3CSK4-primed human monocyte-derived macrophages (hMDMs) were transfected with the indicated concentration of purified LOS from WT *Yersinia pestis (Yp)* (**A, E**), penta-acylated *Yp* (*Yp*Δ*msbB* and *Yp*Δ*lpxP*) (**B, F, and C, G**), and tetra-acylated *Yp* (*Yp*Δ*msbB/lpxP*) (**D, H**). After transfection with the respective LOS variants for 20 hours, cell death was assessed by LDH release (**A-D**) and IL-1β secretion was assessed by ELISA (**E-H**). Comparing cell death **(I)** and IL-1β release **(J)** of different LOS variants transfected at 2μg/ml. Data are represented as the mean ± SD of triplicate wells from 3-5 different human donors. Data were analyzed by ANOVA followed by Holm-Šídák’s multiple comparisons test, ****P<0.0001, ***P<0.001, **P<0.01, *P<0.05.

These results indicate that in contrast to human TLR4, the human noncanonical inflammasome is robustly activated in response to penta-acylated LOS containing a secondary acyl-chain at the 2’ position or at the 3’ position similarly to hexa-acylated LOS. Furthermore, our data indicate that in contrast to the murine noncanonical inflammasome, tetra-acylated lipid A can activate the human noncanonical inflammasome. However, tetra-acylated lipid A is significantly less stimulatory compared to penta-acylated or hexa-acylated lipid A, suggesting that the presence of either the 12-C or 16:1-C secondary acyl-chain in *Yp* LOS is required to elicit a robust human noncanonical inflammasome response.

### The human noncanonical inflammasome is activated by Y. pestis LOS variants regardless of acyl chain length or phosphorylation state

We next asked whether longer secondary acyl chains within LOS can affect noncanonical inflammasome activation. To investigate this, we utilized a hexa-acylated *Yp* LOS variant containing an additional secondary 16-C acyl-chain that was isolated from *Yp*Δ*msbB* expressing the acyltransferase enzyme PagP (*Yp*Δ*msbB/pagP*^+^)[16, 17]. Transfection of this LOS variant into hMDMs led to similar levels of cell death (**Figure 3A**) and IL-1β secretion (**Figure 3B**) as compared to hMDMs transfected with the WT *Yp* LOS. We then tested whether the phosphorylation state of LOS along with changes in acyl chain position can differentially activate the human noncanonical inflammasome. We utilized a hexa-acylated *Yp* LOS variant which is mono-phosphorylated due to expression of the *Francisella novicida* lipid A C-4’ phosphatase LpxF and also has an added secondary 16-C acyl-chain (*YplpxF*^+^*pagP*^+^)[18]. There were similar levels of cell death and IL-1β secretion after transfection of *YplpxF*^+^/*pagP*^+^ LOS into hMDMs compared to hMDMs transfected with WT *Yp* LOS (**Figure 3A-B**). Furthermore, we observed similar levels of cell death and IL-1β secretion in hMDMs transfected with hexaacylated *YplpxF*^+^ LOS lacking the 4’ phosphate group as compared to WT *Yp* (**Figure 3A-B**). Collectively, these data indicate that the human noncanonical inflammasome can be activated by a wide range of tetra-, penta-, and hexa-acylated *Yp* LOS variants regardless of their lipid A phosphorylation or acylation state.

**Figure 3.**
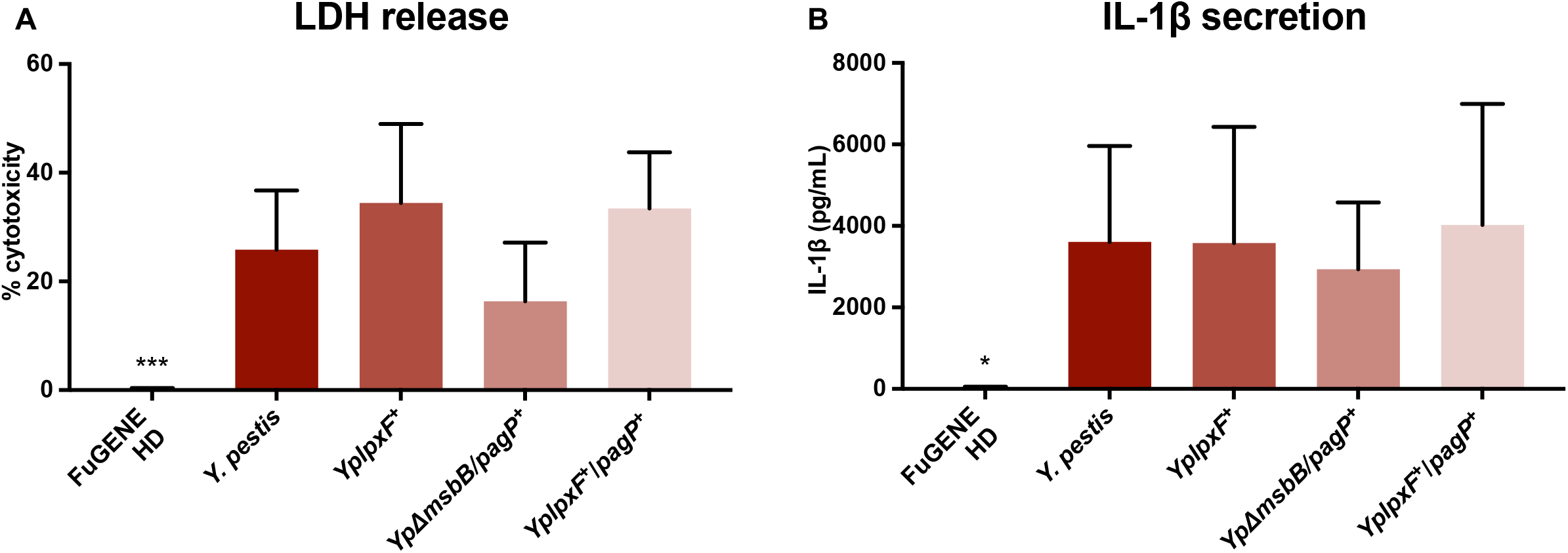
LOS robustly activates the human noncanonical inflammasome regardless of lipid A phosphorylation or acyl chain length. Pam3CSK4-primed hMDMs were transfected with 2 μg/mL of purified LOS from WT *Yp, YplpxF*^+^, *Yp*Δ*msbB/pagP*^+^, *YplpxF*^+^/*pagP*^+^. 20 hours post transfection, cell death was measured by assessing LDH release (**A**) and IL-1β release was measured by ELISA (**B**). Data are represented as the mean ± SD of triplicate wells from seven different healthy human donors. Data were analyzed by ANOVA followed by Holm-Šídák’s multiple comparisons test, ***P<0.001, *P<0.05.

We next asked whether CASP4 and/or CASP5 contribute to detection of these lipid A variants. We used siRNA to silence *CASP4* and/or *CASP5* in primary hMDMs. We observed that knocking down *CASP4* (67% average knockdown), *CASP5* (42% average knockdown), or both (63% and 15% average knockdown for *CASP4* and *CASP5*, respectively) did not significantly affect cell death after transfecting cells with hexa-acylated WT *Yp* LOS or *YplpxF*^+^ LOS, pentaacylated *Yp*Δ*msbB* LOS, or tetra-acylated *Yp*Δ*msbB/lpxP* LOS as compared to control siRNA treated cells (**Figure 4A**). Similarly, we observed that knocking down either *CASP4* or *CASP5* alone did not lead to a significant reduction in IL-1β secretion after transfection with WT *Yp* LOS, *Yp*Δ*msbB* LOS, or *Yp*Δ*msbB/lpxP* LOS (**Figure 4B**). We did notice that individual knockdown of *CASP4* or *CASP5* led to a significant decrease in IL-1β release in hMDMs transfected with *YplpxF*^+^ LOS, which is the hexa-acylated LOS that is missing a 4’ phosphate group. However, knocking down both *CASP4* and *CASP5* significantly decreased IL-1β release after transfecting cells with hexa-acylated WT *Yp* LOS or *YplpxF*^+^ LOS, as well as pentaacylated *Yp*Δ*msbB* LOS compared to control siRNA treated cells (**Figure 4B**). These data suggest that CASP4 and CASP5 both contribute to recognizing LOS containing 3’ and 2’ O-linked acyl chains, which are absent from tetra-acylated *Yp*Δ*msbB/lpxP* LOS. In addition, these data indicate that the robust IL-1β release seen after transfecting hMDMs with the different LOS variants relies on both CASP4 and CASP5, suggesting that one caspase can compensate for the absence of the other to mediate noncanonical inflammasome activation.

**Figure 4.**
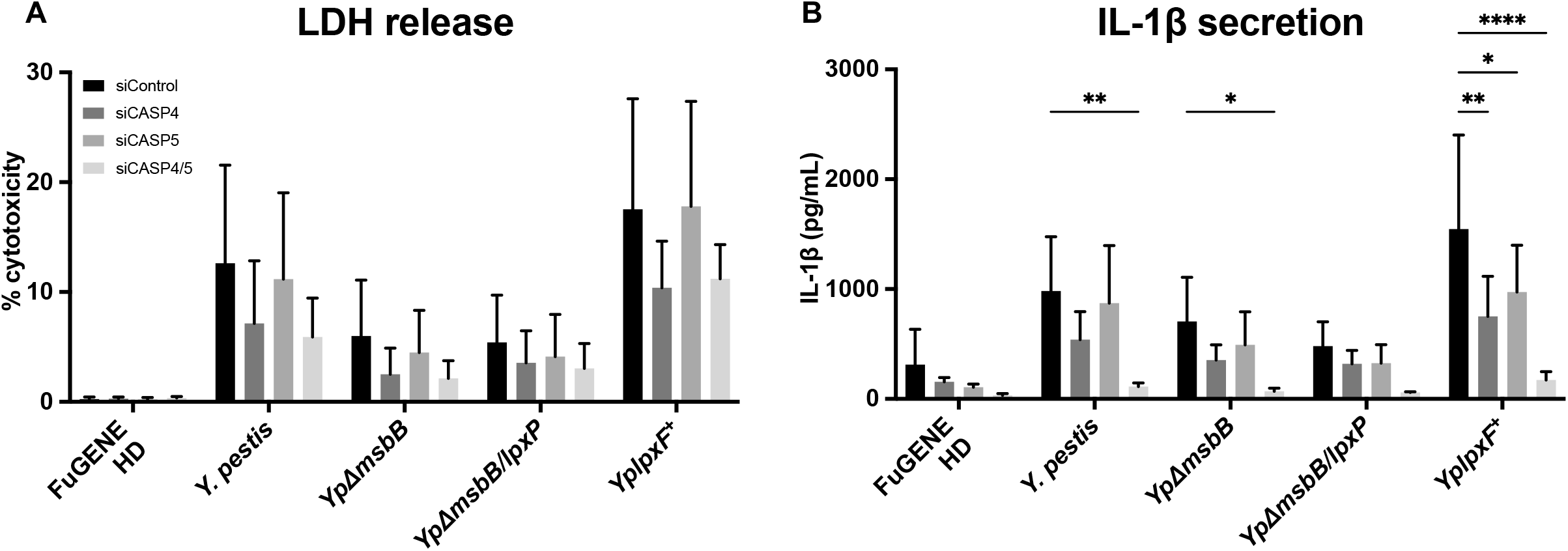
Both caspase-4 and caspase-5 are necessary for maximal inflammasome response to LOS variants. Pam3CSK4-primed hMDMs were transfected with control siRNA or. siRNA against *CASP4* and/or *CASP5*. 24 h after siRNA-mediated knockdown, cells were transfected with 2 μg/mL of purified LOS from WT *Yp*, *Yp*Δ*msbB, Yp*Δ*msbB/lpxP*, or *YplpxF*^+^. 20 hours post LOS transfection, cell death was measured by assessing LDH release (**A**) and IL-1β release was measured by ELISA (**B**). Data are represented as the mean ± SD of triplicate wells from four different healthy human donors. Data were analyzed by ANOVA followed by Holm-Šídák’s multiple comparisons test, ****P<0.0001, **P<0.01, *P<0.05.

### Core oligosaccharide is required for maximum noncanonical inflammasome activation by LOS in human macrophages

Lipid A is sufficient for human TLR4 activation, and absence of the core oligosaccharide does not reduce human or mouse TLR4 activation in response to lipid A when compared to LOS (**Supplemental Figure 2 and** [14]). Previous studies indicate that lipid A is sufficient for noncanonical inflammasome activation in both human and mouse macrophages [3–5], and affinity measurements indicate similar binding of LPS and lipid A to CASP4 [6], suggesting that lipid A and LPS might induce similar levels of noncanonical inflammasome activation. However, it is unknown whether the presence or absence of the core oligosaccharide affects human noncanonical inflammasome responses to lipid A. To address this question, we compared the noncanonical inflammasome response to purified lipid A lacking the core oligosaccharide or lipooligosaccharide (LOS) derived from each of the seven LOS variants. Surprisingly and in contrast to the ability of these identical lipid A preparations to stimulate the murine noncanonical inflammasome [14], purified lipid A variants induced significantly decreased LDH (**Figure 5A**) and IL-1β release (**Figure 5B**) as compared to their respective LOS variants, suggesting that the core oligosaccharide is important for stimulating human noncanonical inflammasome responses. Using an alternate route of lipid A delivery via co-administration with the bacterial pathogen *Listeria monocytogenes* [4], which uses the pore-forming toxin listeriolysin O to escape from the phagosome into the host cell cytosol, also led to decreased responses to lipid A relative to their corresponding LOS structures **(Supplemental Figure 3)**. Taken together, these data indicate that the core oligosaccharide is necessary for maximal human noncanonical inflammasome responses to *Y. pestis* LOS.

**Figure 5.**
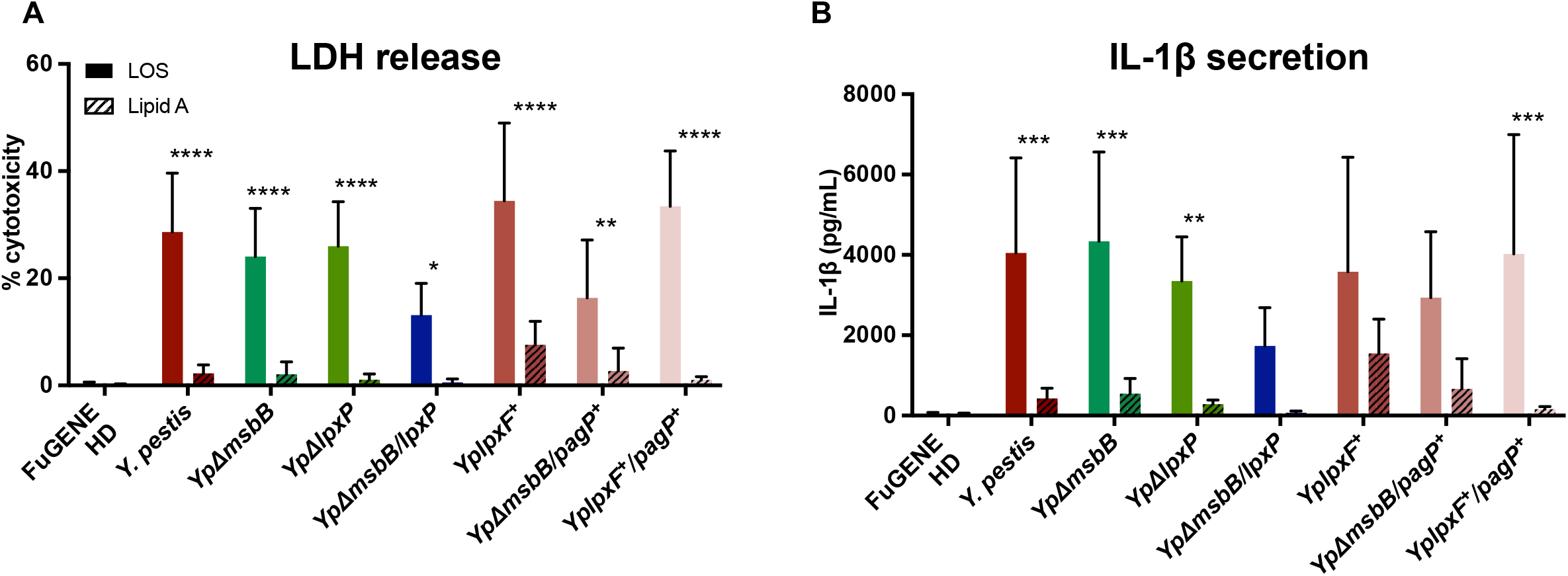
LOS core oligosaccharide is important for maximal human inflammasome responses. Pam3CSK4-primed hMDMs were transfected with 2 μg/mL of purified lipid A or LOS from WT *Yp, Yp*Δ*msbB, Yp*Δ*lpxP, Yp*Δ*msbB/lpxP, YplpxF*^+^, *Yp*Δ*msbB/pagP*^+^, *YplpxF*^+^/*pagP*^+^. After transfection with the respective variants for 20 hours, cell death was measured by assessing LDH release (**A**) and IL-1β secretion (**B**) and lipid A-treated cells (striped bars) were compared to LOS-treated cells (solid bars). Data are represented as the mean ± SD of triplicate wells from 5-9 different human donors. Data were analyzed by ANOVA followed by Holm-Šídák’s multiple comparisons test, ****P<0.0001, ***P<0.001, **P<0.01, *P<0.05.

### Expression of Caspase-4 in murine macrophages confers responsiveness to hypo-acylated LOS and lipid A

Previous studies and our data demonstrate that tetra-acylated lipid A fails to activate murine Casp11 but can activate human CASP4 [3–5, 14]. These findings raise the question of whether the ability of human macrophages to respond to tetra-acylated lipid A is due to a property intrinsic to CASP4 itself or another feature unique to human macrophages. To distinguish between these possibilities, we used *Casp11*^-/-^*Casp4^Tg^* mice, which lack the murine *Casp11* ortholog, but express human *CASP4* as a transgene [19]. We found that expression of the *CASP4* transgene restored responsiveness of *Casp11*^-/-^ BMDMs to hexa-acylated LOS from WT *Yp*, as well as to both penta-acylated LOS species from *Yp*Δ*msbB* and *Yp*Δ*lpxP* (**Supplemental Figure 4**). *Casp11*^-/-^*Casp4^Tg^* BMDMs were also able to respond to tetra-acylated LOS *Yp*Δ*msbB/lpxP*. This contrasted with WT BMDMs expressing endogenous *Casp11*, as they did not respond to *Yp*Δ*msbB/lpxP* tetra-acylated LOS or *Yp*Δ*msbB* penta-acylated LOS (**Supplemental Figure 4** and [14]). These data indicate that the ability of the human noncanonical inflammasome to respond to a range of *Yp* LOS variants regardless of their acylation state is a property intrinsic to CASP4.

Differences in the ability of the human noncanonical inflammasome to respond to lipid A relative to LOS could also potentially be due to an intrinsic property of CASP4, or a caspase-independent difference between human and murine macrophages. To distinguish between these possibilities, we delivered purified lipid A from the different LOS structure variants into wild type, *Casp11*^-/-^, and *Casp11*^-/-^*Casp4^Tg^* BMDMs. As expected, there were no differences in cell death or IL-1β release between wild type BMDMs transfected with LOS or lipid A (**Figure 6A-B**), and this LDH and IL-1β release were dependent on Casp11 (**Figure 6C-D**)[14]. *Casp11*^-/-^ *Casp4^Tg^* BMDMs showed equivalent levels of LDH and IL-1β release in response to transfected LOS and lipid A (**Figure 6E-F, Supplemental Figure 4**).

**Figure 6.**
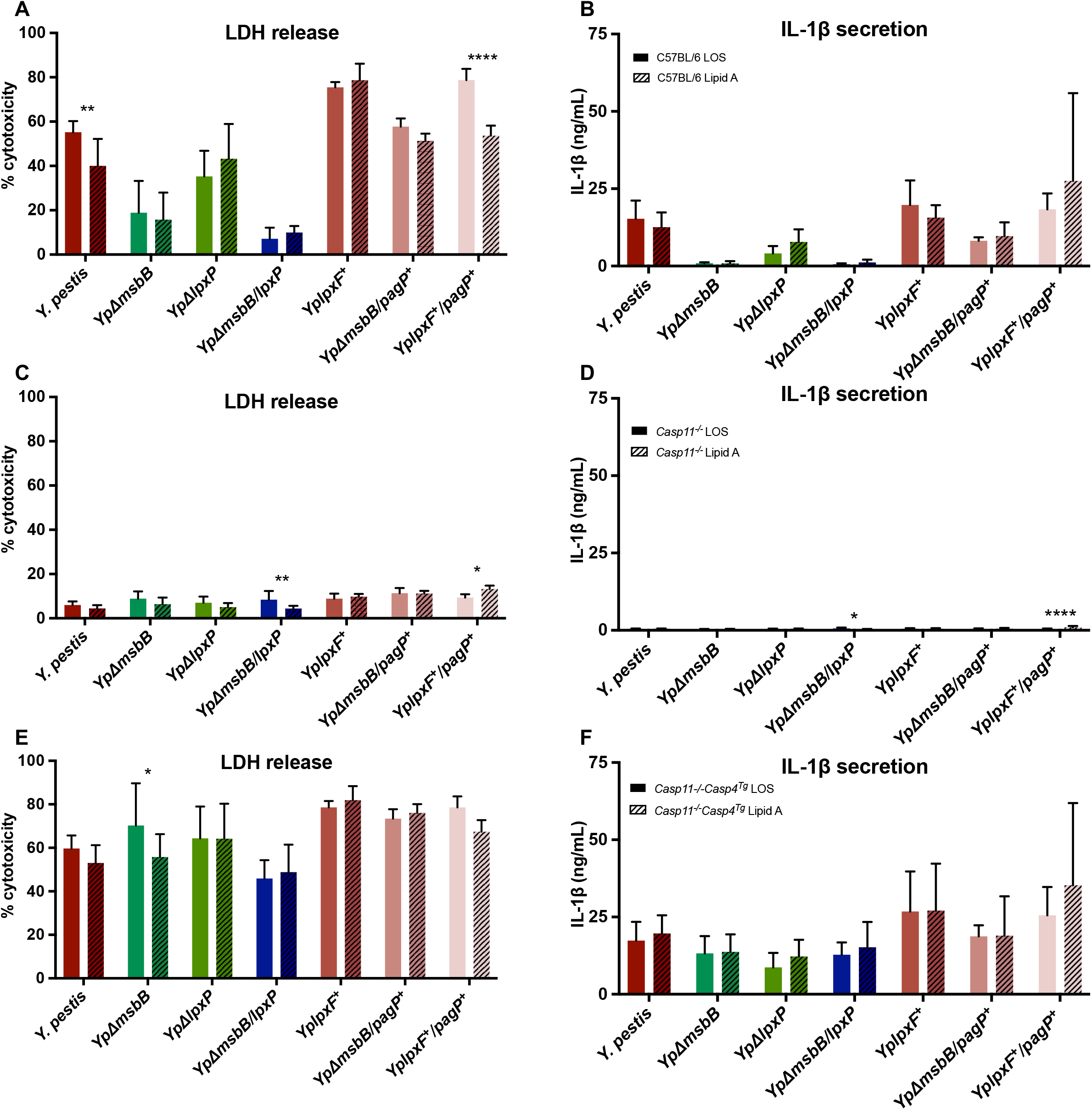
Ectopic expression of caspase-4 confers onto mouse macrophages the ability to respond to both LOS and lipid A. Pam3CSK4-primed BMDMs from wild type (C57BL/6) (**A-B**), *Casp11*^-/-^ (**C-D**), and *Casp11*^-/-^ *Casp4^Tg^* (**E-F**) mice were transfected with the 2 μg/ml of LOS (solid bars) or lipid A (striped bars) from WT *Yp*, penta-acylated *Yp* (*Yp*Δ*msbB* and *Yp*Δ*lpxP*), tetra-acylated *Yp* (*Yp*Δ*msbB/lpxP*), and hexa-acylated *Yp* (*YplpxF*^+^, *Yp*Δ*msbB/pagP*^+^, *YplpxF*^+^/*pagP*^+^). After transfection with the respective LOS or lipid A variants for 20 hours, cell death was measured by LDH release (**A, C, and E**) and IL-1β secretion was measured by ELISA (**B, D, and F**). Data are mean ± SD of three independent experiments performed in triplicate. Data were analyzed by ANOVA followed by Holm-Šídák’s multiple comparisons test, ****P<0.0001, **P<0.01, *P<0.05.

These findings indicate that human macrophages require core oligosaccharide to respond robustly to cytosolic LOS, whereas mouse macrophages do not, suggesting the presence of additional accessory factors in murine cells that facilitate noncanonical inflammasome assembly or activation in response to lipid A.

## Discussion

Defining the minimum stimulatory components of LPS/LOS/lipid A that are required to activate human TLR4 and CASP4/5 is critical to allow for the advance of vaccine adjuvants and immunomodulatory therapeutics, as well as more effective treatment of pathogenic Gramnegative bacterial infections. In this study, we used the BECC process to create a series of *Yersinia pestis*-derived LOS with distinct lipid A structures that contain or lack specific acyl-chains and/or phosphoryl groups and used these LOS structures to interrogate structure-activity relationships between LOS and TLR4, as well as the CASP4/5 noncanonical inflammasome.

Our results show that human TLR4 is activated in response to hexa-acylated LOS and penta-acylated LOS specifically containing the 2’ secondary 16:1-C acyl chain, but not tetraacylated LOS or penta-acylated LOS missing the 2’ secondary acyl chain (**Figure 1**). In contrast, CASP4/5 responded to stimulation by hexa-, penta-, and tetra-acylated LOS, with tetra-acylated eliciting a lower response (**Figures 2 and 3**). Furthermore, CASP4/5 responded to LOS irrespective of lipid A acyl chain length. Recently, a study by Gauthier *et al*. found that deep-sea microbes from the *Moritella* genus maintain basic LPS structures but many are not able to act as effective TLR4 ligands, which was attributed to differences in LPS acyl-chain length [20].

The contribution of phosphate groups to the strength of innate immune response activated by LPS molecules was also of interest to us, as its potential applicability is highlighted by the use of mono-phosphoryl lipid A (MPL) as an effective vaccine adjuvant that stimulates immune responses without inducing pathogenic inflammation [21–23]. Although removing a phosphate is attributed to attenuation of signaling through TLR4, recently published studies indicate that with structural changes to the acyl-chain length and arrangement of lipid A, signal strength can be attenuated even if LPS is bis-phosphorylated [24–26]. Intriguingly, we found that using our hexaacylated LOS structure, we found that both TLR4 and the noncanonical inflammasome were activated in response to mono- and bis-phosphorylated hexa-acylated LOS (**Figures 1 and 3**), indicating that changes in LOS phosphorylation state have no effect on activation of these innate immune pathways.

An unexpected finding from our study is the importance of the core oligosaccharide for maximal human noncanonical inflammasome activation (**Figure 5)**, whereas the core is not required for noncanonical inflammasome responses in mouse macrophages. Although previous studies indicated that presence of the core oligosaccharide allows for maximum TLR4 stimulation [27–29], we found that our lipid A lacking the core oligosaccharide did not attenuate TLR4 activation. We then investigated whether the ability of human macrophages to broadly recognize LOS variants in a core oligosaccharide-dependent manner was intrinsic to human CASP4 or another feature unique to human macrophages. Using mouse *Casp11*^-/-^ macrophages expressing a human *CASP4* transgene, we found that CASP4 confers onto mouse BMDMs the ability to recognize different LOS variants regardless of acylation state. These data indicate that the ability of human macrophages to respond broadly to different LOS variants is a property intrinsic to CASP4. Interestingly, we found that in contrast to human macrophages, mouse BMDMs similarly responded robustly to both LOS and lipid A when expressing CASP4 (**Figure 6**). These data indicate that the ability of mouse BMDMs to respond equally to both LOS and lipid A, as compared to the differential response observed in hMDMs, could be due to the presence of additional mouse-specific host factors in BMDMs that are not present in hMDMs or vice versa.

One host factor that contributes to intracellular LPS recognition are guanylate binding proteins (GBPs), which are a subfamily of interferon-inducible GTPases involved in cell-autonomous immune responses against intracellular bacterial pathogens [30, 31]. It is possible that differences in GBP composition or function are responsible for the differential ability of humans and mice to respond to LOS versus lipid A. Notably, mice have 11 GBPs whereas humans only have 7 GBPs, and GBPs may also exhibit species-specific functional differences [32]. In particular, murine GBPs are thought to promote inflammasome responses by mediating lysis of cytosolic bacteria, promoting rupture of pathogen-containing vacuoles, and by binding LPS or outer membrane vesicles [33–36]. Human GBPs have been shown to inhibit bacterial spread, bind LPS, disrupt the bacterial cell envelope, and control inflammasome assembly at the surface of cytosolic bacteria [37–43]. Although the mouse GBPs encoded on chromosome 3 do not contribute significantly to Casp11 activation by *Y. pestis* LOS in murine systems [14], the human orthologs may play a role. Future studies will be needed to investigate GBPs or other host factors that contribute to activation of the murine and human noncanonical inflammasome.

Our data show that the human noncanonical inflammasome is promiscuous, as it responded to all of the tetra-, penta-, and hexa-acylated *Y. pestis* LOS variants tested. In contrast, Harberts, Grubaugh *et al*. (companion manuscript) show that the mouse noncanonical inflammasome is more selective, as it failed to respond to tetra-acylated lipid A, and only responded to one of the penta-acylated variants in addition to the hexa-acylated variants [14]. Mouse BMDMs expressing human CASP4 were also able to respond to all of the LOS variants. However, we found that although hMDMs mount inflammasome responses to tetra-acylated LOS, the response was significantly lower compared to what was observed with penta- or hexaacylated LOS. Although our studies were conducted in the absence of IFNγ priming, our findings are largely in agreement with a previous study that employed IFNγ priming and showed that human CASP4 can be activated by both tetra- and hexa-acylated LPS [5]. The mechanisms underlying the broad recognition of these LOS variants in humans is unclear but could be due to differences in the sequence and resulting structure of human CASP4 and CASP5 compared to mouse Casp11.

In addition, the human noncanonical inflammasome consists of two lipid A sensors, CASP4 and CASP5. Previous studies indicate that CASP4 plays a more dominant role in allowing human macrophages to respond to LPS [6, 44, 45]. We show that both CASP4 and CASP5 contribute to detection of some of the LOS variants, as we observed significantly decreased IL-1β release when both CASP4 and CASP5 were knocked down (**Figure 4**). This finding begins to address the necessity for CASP4 and CASP5 in the detection of these LOS variants. Additional studies are required to fully parse out whether there are differences in how CASP4 and CASP5 respond to LOS variants.

Taken together, our findings reveal that human TLR4 and CASP4/5 differ in their response to tetra-acylated LOS variants and requirement for the presence of the core oligosaccharide. Given the capacity of bacterial pathogens to generate a wide variety of lipid A structures that have the potential to evade either TLR4 or CASP4/5, our findings suggest that the ability of TLR4 and CASP4/5 to detect both overlapping and distinct LOS structures enables the innate immune system to sense a wider range of lipid A structures than would be possible with either sensor on its own, thereby imposing a constraint on the ability of pathogens to evolve LPS or LOS structures that can evade immune detection. Furthermore, our data indicate that in contrast to mice, the human noncanonical inflammasome can respond to a broader range of LOS variants [14], indicating that humans may mount immune responses against a broader range of Gram-negative bacterial pathogens. Collectively, these findings provide a foundation for further understanding the mechanisms underlying these species-specific differences. Additional evaluation of LOS or LPS structures from a wide range of bacterial species will help broaden our understanding of TLR4 and CASP4/5 recognition and provide insight into host:pathogen interactions. In addition, our study indicates that future studies of human TLR4 and CASP4/5 to enable selective targeting of human TLR4 and the noncanonical inflammasome as a therapeutic approach against Gram-negative sepsis.

## Acknowledgements

We thank members of the Shin, Ernst, and Brodsky laboratories for helpful scientific discussions. We thank the Human Immunology Core of the Penn Center for AIDS Research and Abramson Cancer Center for providing purified primary human monocytes. This work is supported in part by R01AI118861 (SS), R01AI123243 (SS), HHS-NIH-NIAID-BAA2017 (RKE), AI128530 (IEB), AI139102A1 (IEB), K12GM081259 (JAF), T32AI095190 (EMH), T32AR076951 (DG), National Science Foundation Graduate Fellowship DGE-1845298 (to A.R.B.), and a Burroughs-Welcome Fund Investigators in the Pathogenesis of Infectious Diseases Award (SS and IEB).

## Materials and Methods

### Ethics Statement

All studies on primary human monocyte-derived macrophages (hMDMs) were performed in compliance with the requirements of the US Department of Health and Human Services and the principles expressed in the Declaration of Helsinki. Samples obtained from the University of Pennsylvania Human Immunology Core are considered to be a secondary use of deidentified human specimens and are exempt via Title 55 Part 46, Subpart A of 46.101 (b) of the Code of Federal Regulations. All animal studies were performed in compliance with the federal regulations set forth in the Animal Welfare Act [10], the recommendations in the Guide for the Care and Use of Laboratory Animals of the National Institutes of Health, and the guidelines of the University of Pennsylvania Institutional Animal Use and Care Committee. All protocols used in this study were approved by the Institutional Animal Care and Use Committee at the University of Pennsylvania (protocols #804523 and #804928).

### Mice

Bone marrow-derived macrophages from WT C57BL/6 (Jackson Laboratory), *Casp11*^-/-^ (Jackson Laboratory)[46], and *Casp11*^-/-^*Casp4^Tg^* (from Joseph Buxbaum laboratory)[19] mice were used in this study.

### Lipooligosaccharide (LOS) and Lipid A Variants

LOS structural variants were created using bacterial enzyme combinatorial chemistry in the *Yersinia pestis* (*Yp*) KIM6+ strain, an avirulent, non-select agent variant [47]. Due to a mutation that only allows for the addition of one O-antigen unit, *Yp* makes only LOS, not LPS. Lipid A structural variants include *Yp*Δ*msbB, Yp*Δ*lpxP, Yp*Δ*msbB/lpxP, YplpxF*^+^, *YplpxF*^+^/*pagP*^+^, and *Yp*Δ*msbB/pagP*^+^ [13, 15, 17] (**Supplemental Figure 1**). Briefly, to the tetra-acylated lipid A structure: *msbB* adds a 3’ O-linked 12C acyl chain, *lpxP* adds a 2’ O-linked 16:1C acyl chain, and *lpxF* removes the 4’ phosphate. Bacteria were cultured, harvested, and LOS extracted as described in [14]. Lipid A was collected from the extracted LOS using a mild acid hydrolysis as previously described [13].

### Cell Culture

Reporter cell lines with a SEAP gene under control of the NFκB promoter were used to determine TLR4 activation levels in response to varied LOS structures. These cells, HEK-Blue hTLR4 cells (InvivoGen) were cultured in DMEM and THP-1 Dual cells (InvivoGen) were cultured in RPMI, both media were supplemented with 10% (vol/vol) heat-inactivated FBS, 2 mM L-glutamine, 100 IU/mL penicillin, and 100 μg/mL streptomycin. Cells were plated at a density of 100,000 cells/well in a 96-well flat bottom plate and cultures in a 5% CO_2_ humidified incubator. THP-1 Dual cells were cultured with 100 ng/mL Vitamin D_3_ to promote adherence and surface expression of CD14, a TLR4 co-receptor. Cells were stimulated for 18 hours with a 5-log dose range of agonist as previously described [13]. SEAP amounts were then measured in the cell culture supernatants using Quanti-Blue Detection Media (InvivoGen) according to manufacturer’s instructions. The OD_620_ represents the activation level of NF-κB in each well.

Primary human monocytes from deidentified healthy human donors were obtained from the University of Pennsylvania Human Immunology Core. Monocytes were cultured in RPMI supplemented with 10% (vol/vol) heat-inactivated FBS, 2 mM L-glutamine, 100 IU/mL penicillin, 100 μg/mL streptomycin, and 50 ng/mL recombinant human M-CSF (Gemini Bio Products). Cells were cultured for 4 days in 10 mL of media in 10 cm-dishes at 4-5 × 10^5^ cells/mL, followed by addition of 10 mL of fresh growth media for an additional 2 days for complete differentiation into macrophages. The day before macrophage stimulation, cells were rinsed with cold PBS, gently detached with trypsin-EDTA (0.05%) and replated in media without antibiotics and with 25 ng/mL M-CSF in a 48-well plate at a concentration of 1 × 10^5^ cells per well.

Mouse bone marrow-derived macrophages (BMDMs) were cultured in DMEM supplemented with 10% fetal bovine serum, 10mM HEPES, 1mM sodium pyruvate, and 30% L929 cell conditioned medium. Cells were grown for 6-7 days in non-tissue culture treated plates before reseeding into 48-well tissue culture plates at a density of 1 × 10^5^ cells per well 20 hours before LOS and lipid A transfection.

### LOS and Lipid A Intracellular Delivery

Primary human monocyte derived macrophages (hMDMs) or murine bone marrow-derived macrophages (BMDMs) were primed with 1 μg/mL or 400 ng/mL Pam3CSK4 (InvivoGen), respectively for 4 h. The media was then replaced with 300 μL of serum-free Opti-MEM Reduced Serum Medium (Thermo Fisher Scientific) per well, and hMDMs were either mock-transfected with FuGENE HD (Promega) alone or treated with a mixture of 0.75 μL FuGENE HD [0.25% (vol/vol)] plus LOS or lipid A (2 μg/mL or at the indicated concentrations). Plates were then centrifuged at 805 × g for 5 min before culturing at 37 °C for 20 h. *Listeria monocytogenes* co-infection of lipid A into hMDMs was performed as in [4]. Briefly, primary hMDMs were primed with Pam3CSK4 for 4 hours then infected with *L. monocytogenes* strain 10403S at a MOI of 5 in the presence of 2 μg/ml lipid A. After one hour of infection, 20 μg/mL gentamicin was added to kill extracellular bacteria. Four hours after infection, cell supernatants were harvested to assess cell death and cytokine secretion.

### LDH Cytotoxicity Assay

Primary hMDMs and BMDMs were transfected in a 48-well plate as described above and harvested supernatants were assayed for cell death by measuring loss of cellular membrane integrity via lactate dehydrogenase (LDH) activity. LDH release was quantified using an LDH Cytotoxicity Detection Kit (Takara BioProduct) according to the manufacturer’s instructions and normalized to mock-infected cells.

### ELISA

To measure human IL-1β, primary hMDMs were transfected in a 48-well plate as described above and harvested supernatants were assayed for cytokine levels using an ELISA kit for human IL-1β (BD Biosciences) according to the manufacturer’s instructions.

To measure mouse IL-1β, supernatants and recombinant cytokine standards were applied to anti-IL-1β antibody-coated (eBioscience) Immulon ELISA plates (ImmunoChemistry Technologies). IL-1β was detected using biotinylated anti IL-1β (eBioscience) and streptavidin conjugated to horseradish peroxidase (BD Biosciences). Peroxidase activity was detected using an o-phenylenediamine hydrochloride (Sigma-Aldrich) solution in citrate buffer. Reactions were stopped with 3M H2SO4 and absorbance at 490nm read with a spectrophotometer.

### siRNA Knockdown

All Silencer Select siRNA oligos were purchased from Ambion (Life Technologies). For CASP4, the siRNA used was siRNA identification s2414, and s2417 for CASP5. To knockdown CASP4 or CASP5 alone, 30 nM of the respective oligo was used per well. To knockdown both CASP4 and CASP5, 15 nM of each oligo was used per well. As a control, Silencer Select negative control siRNAs (Silencer Select Negative Control No. 1 siRNA 4390843 and Silencer Select Negative Control No. 2 siRNA 4390846) were used at 15 nM each per well. Transfection of the pooled siRNAs into macrophages was performed using Lipofectamine RNAiMAX transfection reagent (Thermo Fisher Scientific) following the manufacturer’s protocol. Treatment with appropriate siRNAs was performed for 24 h.

### qRT-PCR Analysis

Cells were lysed, and RNA was isolated using the RNeasy Plus Kit (Qiagen). Synthesis of the first strand cDNA was performed using SuperScript II reverse transcriptase and oligo (dT) primer (Invitrogen). qPCR was performed with the CFX96 real-time system (Bio-Rad) using the SsoFast EvaGreen Supermix with the Low ROX kit (Bio-Rad). The following primers from PrimerBank [48–50] were used. The PrimerBank identifications are *CASP4* (73622124c2), *CASP5* (209870072c1) and *HPRT*(164518913c1; all 5′–3′):

*CASP4* forward: TCTGCGGAACTGTGCATGATG;
*CASP4* reverse: TGTGTGATGAAGATAGAGCCCAT;
*CASP5* forward: TCACCTGCCTGCAAGGAATG;
*CASP5* reverse: TCTTTTCGTCAACCACAGTGTAG;
*HPRT* forward: CCTGGCGTCGTGATTAGTGAT; and
*HPRT* reverse: AGACGTTCAGTCCTGTCCATAA.

For analysis, mRNA levels of siRNA-treated cells were normalized to control siRNA-treated cells using the 2^-ΔΔCT^ (cycle threshold) [51] method to calculate fold induction.

### Statistical Analyses

All graphed data and ANOVA analyses were carried out in GraphPad Prism (San Diego, California). ANOVA was followed by multiple comparison with the Holm-Šídák’s post-test. The resulting significance levels are indicated in the figures. All p-values and significance levels are indicated in the figures and figure legends.

## References

1. Peleg, A.Y. and D.C. Hooper, Hospital-acquired infections due to gram-negative bacteria. N Engl J Med, 2010. 362(19): p. 1804–13.

2. Marshall, J.C., Why have clinical trials in sepsis failed? Trends Mol Med, 2014. 20(4): p. 195–203.

3. Kayagaki, N., et al., Noncanonical inflammasome activation by intracellular LPS independent of TLR4. Science, 2013. 341(6151): p. 1246–9.

4. Hagar, J.A., et al., Cytoplasmic LPS activates caspase-11: implications in TLR4-independent endotoxic shock. Science, 2013. 341(6151): p. 1250–3.

5. Lagrange, B., et al., Human caspase-4 detects tetra-acylated LPS and cytosolic Francisella and functions differently from murine caspase-11. Nat Commun, 2018. 9(1): p. 242.

6. Shi, J., et al., Inflammatory caspases are innate immune receptors for intracellular LPS. Nature, 2014. 514(7521): p. 187–92.

7. Bertani, B. and N. Ruiz, Function and Biogenesis of Lipopolysaccharides. EcoSal Plus, 2018. 8(1).

8. Hajjar, A.M., et al., Human Toll-like receptor 4 recognizes host-specific LPS modifications. Nat Immunol, 2002. 3(4): p. 354–9.

9. Montminy, S.W., et al., Virulence factors of Yersinia pestis are overcome by a strong lipopolysaccharide response. Nat Immunol, 2006. 7(10): p. 1066–73.

10. Matsuura, M., et al., Immunomodulatory effects of Yersinia pestis lipopolysaccharides on human macrophages. Clin Vaccine Immunol, 2010. 17(1): p. 49–55.

11. Steeghs, L., et al., Differential activation of human and mouse Toll-like receptor 4 by the adjuvant candidate LpxL1 of Neisseria meningitidis. Infect Immun, 2008. 76(8): p. 3801–7.

12. Reeves, P.R., E. Pacinelli, and L. Wang, O antigen gene clusters of Yersinia pseudotuberculosis. Adv Exp Med Biol, 2003. 529: p. 199–206.

13. Gregg, K.A., et al., Rationally Designed TLR4 Ligands for Vaccine Adjuvant Discovery. mBio, 2017. 8(3).

14. Harberts, E., et al., Position-specific secondary acylation determines detection of lipid A by murine TLR4 and caspase-11. The Journal of Immunology, 2021.

15. Rebeil, R., et al., Characterization of late acyltransferase genes of Yersinia pestis and their role in temperature-dependent lipid A variation. J Bacteriol, 2006. 188(4): p. 1381–8.

16. Correction for Chandler et al., Early evolutionary loss of the lipid A modifying enzyme PagP resulting in innate immune evasion in Yersinia pestis. Proc Natl Acad Sci U S A, 2020. 117(51): p. 32817.

17. Chandler, C.E., et al., Early evolutionary loss of the lipid A modifying enzyme PagP resulting in innate immune evasion in Yersinia pestis. Proc Natl Acad Sci U S A, 2020. 117(37): p. 22984–22991.

18. Jones, J.W., et al., Comprehensive structure characterization of lipid A extracted from Yersinia pestis for determination of its phosphorylation configuration. J Am Soc Mass Spectrom, 2010. 21(5): p. 785–99.

19. Kajiwara, Y., et al., A critical role for human caspase-4 in endotoxin sensitivity. J Immunol, 2014. 193(1): p. 335–43.

20. Gauthier, A.E., et al., Deep-sea microbes as tools to refine the rules of innate immune pattern recognition. Sci Immunol, 2021. 6(57).

21. Qureshi, N., K. Takayama, and E. Ribi, Purification and structural determination of nontoxic lipid A obtained from the lipopolysaccharide of Salmonella typhimurium. J Biol Chem, 1982. 257(19): p. 11808–15.

22. Ribi, E., et al., Lipid A and immunotherapy. Rev Infect Dis, 1984. 6(4): p. 567–72.

23. Tomai, M.A., et al., The adjuvant properties of a nontoxic monophosphoryl lipid A in hyporesponsive and aging mice. J Biol Response Mod, 1987. 6(2): p. 99–107.

24. Gregg, K.A., et al., A lipid A-based TLR4 mimetic effectively adjuvants a Yersinia pestis rF-V1 subunit vaccine in a murine challenge model. Vaccine, 2018. 36(28): p. 4023–4031.

25. Harberts, E., et al., Novel lipid A mimetics, BECC438 and BECC470, act as potent adjuvants in bacterial and viral subunit vaccines. The Journal of Immunology, 2020. 204(1 Supplement): p. 166.14–166.14.

26. Zacharia, A., et al., Optimization of RG1-VLP vaccine performance in mice with novel TLR4 agonists. Vaccine, 2021. 39(2): p. 292–302.

27. Zughaier, S., et al., Hexa-acylation and KDO(2)-glycosylation determine the specific immunostimulatory activity of Neisseria meningitidis lipid A for human monocyte derived dendritic cells. Vaccine, 2006. 24(9): p. 1291–7.

28. Gaekwad, J., et al., Differential induction of innate immune responses by synthetic lipid a derivatives. J Biol Chem, 2010. 285(38): p. 29375–86.

29. Muroi, M. and K. Tanamoto, The polysaccharide portion plays an indispensable role in Salmonella lipopolysaccharide-induced activation of NF-kappaB through human toll-like receptor 4. Infect Immun, 2002. 70(11): p. 6043–7.

30. Vestal, D.J., The guanylate-binding proteins (GBPs): proinflammatory cytokine-induced members of the dynamin superfamily with unique GTPase activity. J Interferon Cytokine Res, 2005. 25(8): p. 435–43.

31. Kim, B.H., et al., IFN-inducible GTPases in host cell defense. Cell Host Microbe, 2012. 12(4): p. 432–44.

32. Olszewski, M.A., J. Gray, and D.J. Vestal, In silico genomic analysis of the human and murine guanylate-binding protein (GBP) gene clusters. J Interferon Cytokine Res, 2006. 26(5): p. 328–52.

33. Liu, B.C., et al., Constitutive Interferon Maintains GBP Expression Required for Release of Bacterial Components Upstream of Pyroptosis and Anti-DNA Responses. Cell Rep, 2018. 24(1): p. 155–168 e5.

34. Meunier, E., et al., Caspase-11 activation requires lysis of pathogen-containing vacuoles by IFN-induced GTPases. Nature, 2014. 509(7500): p. 366–70.

35. Meunier, E., et al., Guanylate-binding proteins promote activation of the AIM2 inflammasome during infection with Francisella novicida. Nat Immunol, 2015. 16(5): p. 476–484.

36. Santos, J.C., et al., LPS targets host guanylate-binding proteins to the bacterial outer membrane for non-canonical inflammasome activation. EMBO J, 2018. 37(6).

37. Li, P., et al., Ubiquitination and degradation of GBPs by a Shigella effector to suppress host defence. Nature, 2017.551(7680): p. 378–383.

38. Piro, A.S., et al., Detection of Cytosolic Shigella flexneri via a C-Terminal Triple-Arginine Motif of GBP1 Inhibits Actin-Based Motility. mBio, 2017. 8(6).

39. Wandel, M.P., et al., GBPs Inhibit Motility of Shigella flexneri but Are Targeted for Degradation by the Bacterial Ubiquitin Ligase IpaH9.8. Cell Host Microbe, 2017. 22(4): p. 507–518 e5.

40. Kutsch, M., et al., Direct binding of polymeric GBP1 to LPS disrupts bacterial cell envelope functions. EMBO J, 2020. 39(13): p. e104926.

41. Santos, J.C., et al., Human GBP1 binds LPS to initiate assembly of a caspase-4 activating platform on cytosolic bacteria. Nat Commun, 2020. 11(1): p. 3276.

42. Wandel, M.P., et al., Guanylate-binding proteins convert cytosolic bacteria into caspase-4 signaling platforms. Nat Immunol, 2020. 21(8): p. 880–891.

43. Fisch, D., et al., Human GBP1 is a microbe-specific gatekeeper of macrophage apoptosis and pyroptosis. EMBO J, 2019. 38(13): p. e100926.

44. Casson, C.N., et al., Human caspase-4 mediates noncanonical inflammasome activation against gram-negative bacterial pathogens. Proc Natl Acad Sci U S A, 2015. 112(21): p. 6688–93.

45. Baker, P.J., et al., NLRP3 inflammasome activation downstream of cytoplasmic LPS recognition by both caspase-4 and caspase-5. Eur J Immunol, 2015. 45(10): p. 2918–26.

46. Wang, S., et al., Murine caspase-11, an ICE-interacting protease, is essential for the activation of ICE. Cell, 1998. 92(4): p. 501–9.

47. Sun, W., et al., A live attenuated strain of Yersinia pestis KIM as a vaccine against plague. Vaccine, 2011. 29(16): p. 2986–98.

48. Spandidos, A., et al., A comprehensive collection of experimentally validated primers for Polymerase Chain Reaction quantitation of murine transcript abundance. BMC Genomics, 2008. 9: p. 633.

49. Spandidos, A., et al., PrimerBank: a resource of human and mouse PCR primer pairs for gene expression detection and quantification. Nucleic Acids Res, 2010. 38(Database issue): p. D792–9.

50. Wang, X. and B. Seed, A PCR primer bank for quantitative gene expression analysis. Nucleic Acids Res, 2003. 31(24): p. e154.

51. Livak, K.J. and T.D. Schmittgen, Analysis of relative gene expression data using realtime quantitative PCR and the 2(-Delta Delta C(T)) Method. Methods, 2001. 25(4): p. 402–8.

