## Supplemental Figures for "Human TLR4 and noncanonical inflammasome differ in their ability to respond to distinct lipid A variants"

#### Supplementary Figure Legends

**Supplemental Figure 1. Structures of WT Y*p* and associated variants.**

**Supplemental Figure 2. Core oligosaccharide minimally contributes to hTLR4 stimulatory capacity of structurally varied lipid A molecules.** Lipid A structural variants were cultured with reporter cell lines that overexpress human TLR4/MD-2 (HEK-Blue hTLR4) or express endogenous levels of TLR4 (THP-1 Dual) for 18 hours. Agonists were derived from wild type *E. coli* (red), wild type *Yp* (orange), *Yp*Δ*msbB* (light green), *Yp*Δ*lpxP* (dark green), *Yp*Δ*msbB/lpxP* (blue), *YplpxF*$^{+}$(pink), *YplpxF*$^{+}$/*pagP*$^{+}$ (brown), and *Yp*Δ*msbB/pagP*$^{+}$(purple). Results were graphed using GraphPad Prism v7 with a 4-parameter exponential line of best fit superimposed. Each data point is an average of biological duplicates.

**Supplemental Figure 3. Delivery of lipid A using *Listeria monocytogenes* confirms requirement of core oligosaccharide for maximal inflammasome activation.** Pam3CSK4-primed hMDMs were infected with *L. monocytogenes* in the presence of 2 μg/ml of the different *Yp* lipid A variants for four hours. Subsequently, cell death (**A**) and IL-1β release (**B**) were measured. Solid bars represent lipid A transfected via FuGENE HD and striped bars represent lipid A delivered via. *L. monocytogenes*. Data are represented as the mean ± SD of triplicate wells from three different human donors. Data were analyzed by ANOVA followed by Holm-Šídák's multiple comparisons test, ****P<0.0001.

**Supplemental Figure 4. Caspase-4 is activated in response to LOS and lipid A variants.**

Pam3CSK4-primed BMDMs from wild type (C57BL/6), caspase-11-deficient (*Casp11*$^{-/-}$), and caspase-4 transgenic in caspase-11-deficient (*Casp11*$^{-/-}$*Casp4*$^{Tg}$) mice were transfected with the indicated concentration of LOS or lipid A from WT *Yp* (**A,E**), *Yp*Δ*msbB* (**B,F**), *Yp*Δ*lpxP* (**C, G**), and *Yp*Δ*msbB/lpxP* (**D, H**). After transfection with the respective LOS or lipid A variants for 20 hours, cell death was measured by LDH release (**A-D**) and IL-1β secretion was measured by ELISA (**E-H**). Data are represented as the mean ± SD of three independent experiments performed in triplicate.

**Supplemental Figure 1**

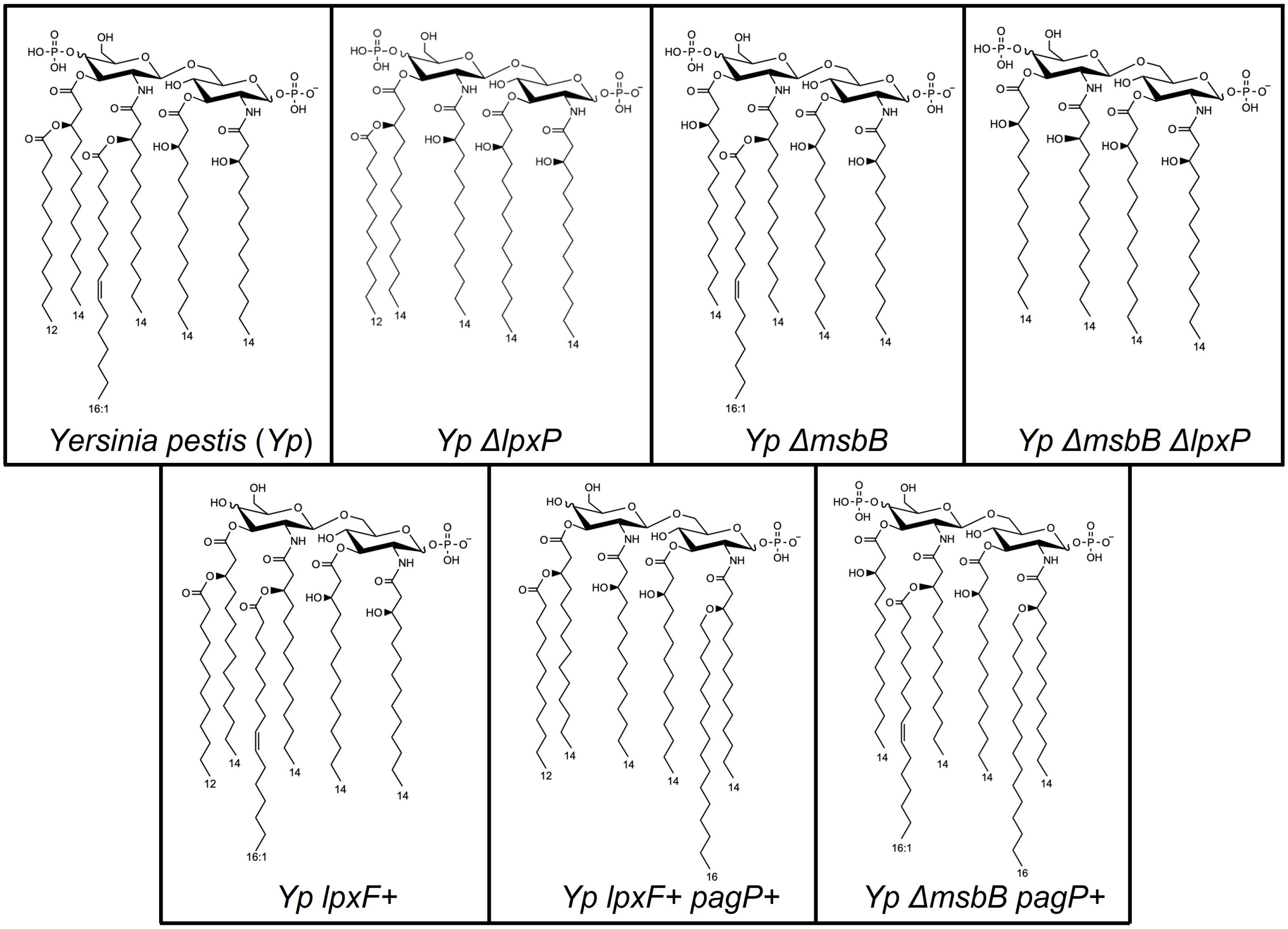

**Supplemental Figure 2**

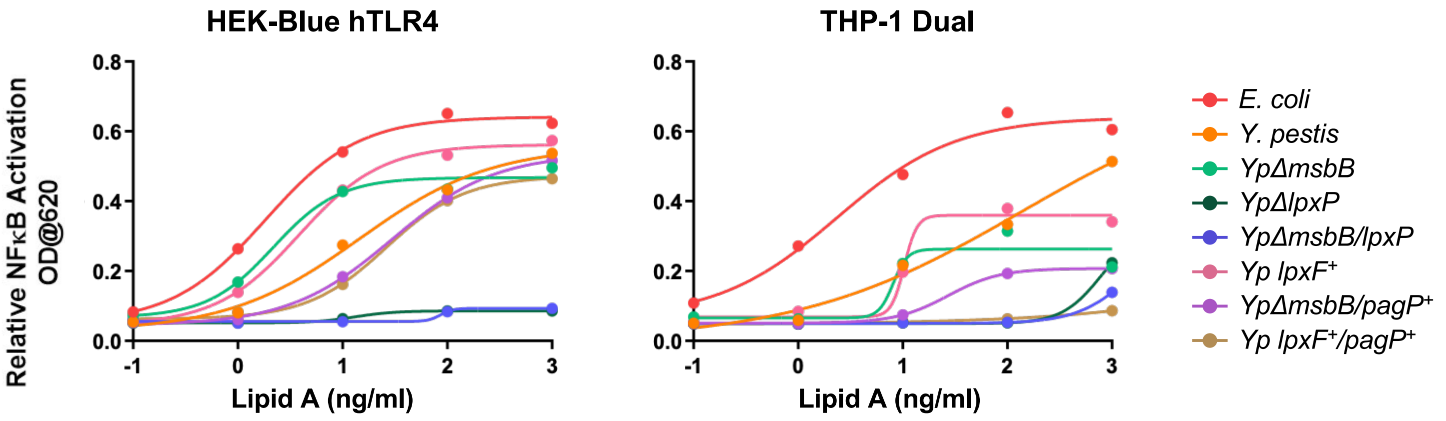

Supplemental Figure 3

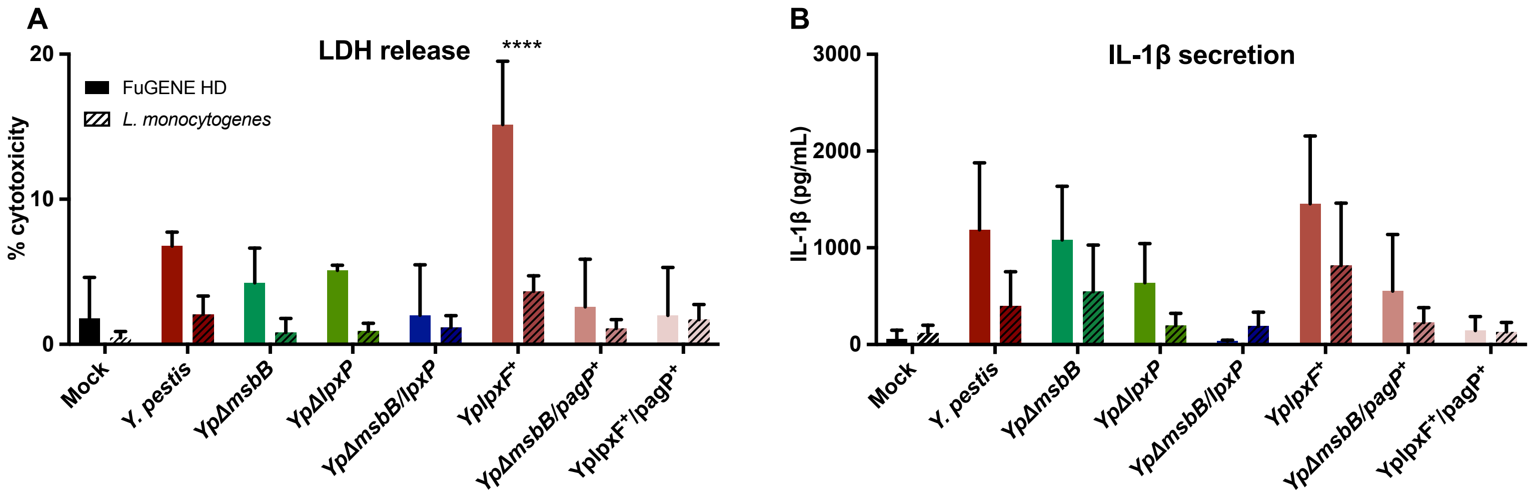

### Supplemental Figure 4

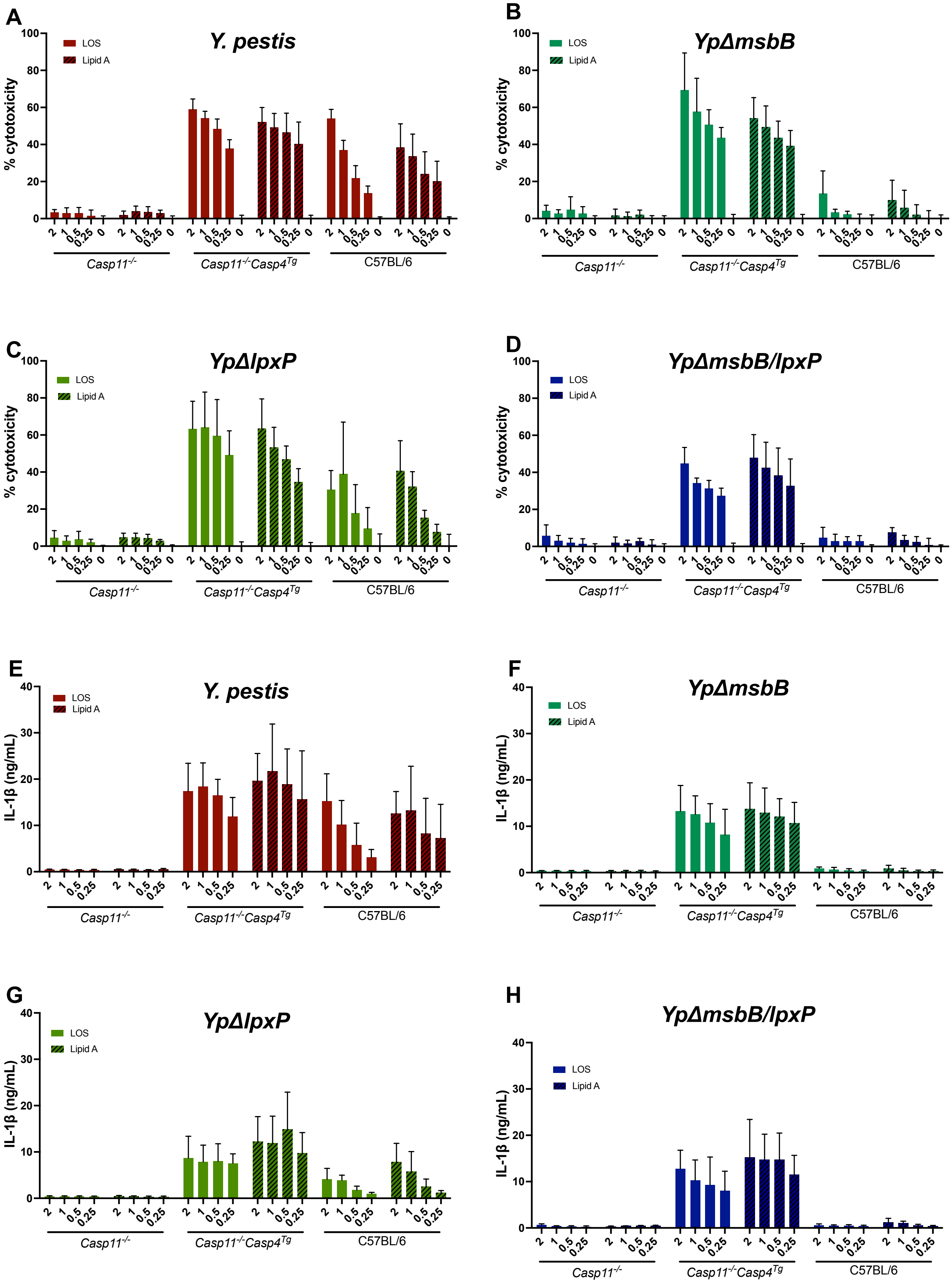
